## Supplementary Notes for "Truly Privacy-Preserving Federated Analytics for Precision Medicine with Multiparty Homomorphic Encryption"

### Supplementary Note 1: Comparison with Existing Works

In Table S1, we compare certain existing solutions and FAMHE on multiple criterion: whether (i) patient levels and (ii) aggregated data are protected, (iii) the data protection satisfies the GDPR definition of anonymity, (iv) the solutions scale with the number of data providers and computing parties, (v) a majority of computing parties can be dishonest (without deviating from the protocol), (vi) the obtained results are the same as if they were computed on a centralized cleartext dataset (i.e., utility), (vii) the system enables multiple types of computations, and (viii) the solution is tested in a real application scenario. A "∼" means that the property is partially fulfilled.

| | Protect. Mecha. | Patient Level Data Protect. | GDPR Anonym. | Intermed. Val. Protect. | Scale w. Parties | Passive Dishonest Majority | Preserve Utility | Comput. Flexibility | Complex Comput. | Appl. To Real Cases |
|---|---|---|---|---|---|---|---|---|---|---|
| AllOfUs[9]<br>Gen. Eng.[10]<br>UKBio.[11] | None | No | No | No | ∼ | No | Yes | Yes | Yes | Yes |
| DataSHIELD[7]<br>Vantage6[8]<br>SHRINE[52]<br>Splink[3]<br>SWARM[4] | Aggregates | ∼ | No | No | Yes | No | ∼ | ∼ | Yes | ∼ |
| Bonomi et al.[16] | DiffP | Yes | No | ∼ | Yes | Yes | No | No | No | ∼ |
| Li et al.[17] | DiffP | Yes | No | ∼ | Yes | Yes | No | No | Yes | No |
| Cho et al.[19] | SMC | Yes | Yes | Yes | No | No | ∼ | No | Yes | Yes |
| Jagadeesh et al. [18] | SMC | Yes | Yes | Yes | No | No | ∼ | No | No | Yes |
| Froelicher et al.[24]<br>Lu et al.[37] | MHE | Yes | Yes | Yes | Yes | Yes | ∼ | ∼ | No | No |
| **FAMHE** | MHE | Yes | Yes | Yes | Yes | Yes | ∼ | ∼ | Yes | Yes |

**Table S1. Comparison of Existing Solutions for biomedical FA.**

### Supplementary Note 2: Secure & Distributed Computation of a Kaplan-Meier Survival Curve

Figure S1 depicts FAMHE secure and federated workflow for the computation of a survival curve (see *Online Methods*). Each $DP_i$ (with $i = 0, \dots, S$) locally computes, encodes and encrypts a vector of the form $n_0^{(i)}, c_0^{(i)}, d_0^{(i)}, \dots, n_T^{(i)}, c_T^{(i)}, d_T^{(i)}$ containing the values $n_j^{(i)}$ (number of survivors), $c_j^{(i)}$ (number of censored), $d_j^{(i)}$ (number of deceased) corresponding to each time point $t_j$ for $t_j = 0, \dots, T$. The DPs' vectors are then collectively aggregated and the final result is collectively switched from the public key $pk$ to the querier's public key, who can decrypt the result with its secret key and generate the curve following Eq. (1) (in *Online Methods*).

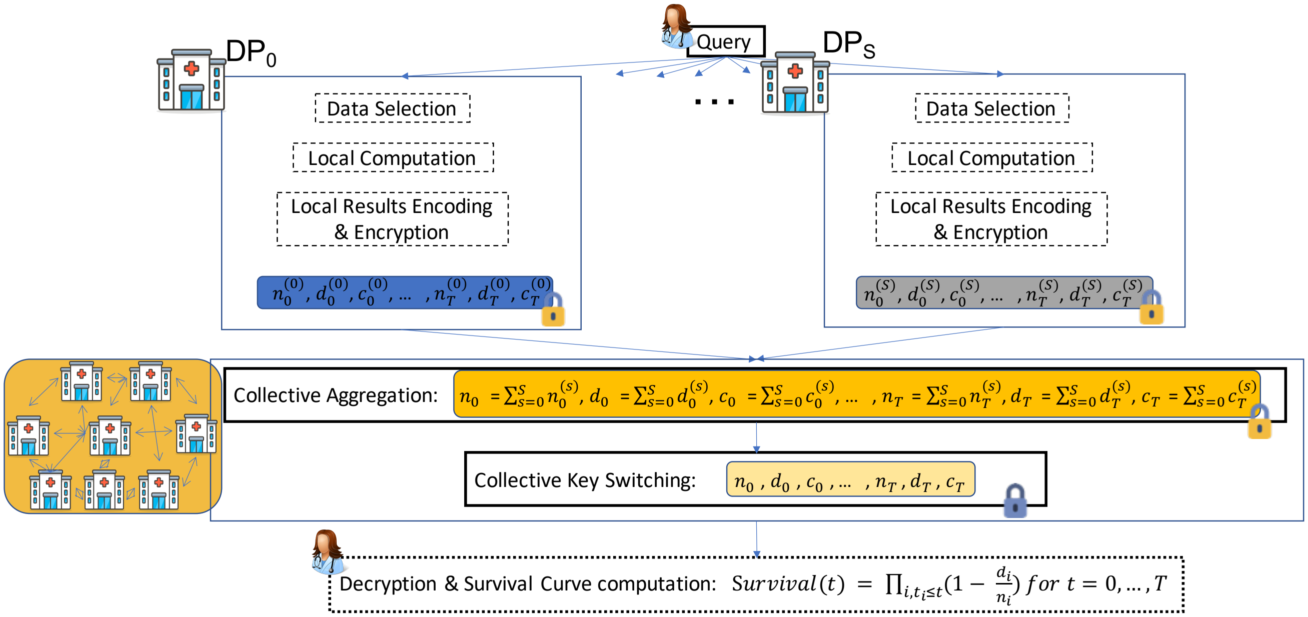


**Figure S1. Secure & Distributed Computation of a Survival Curve.**

### Supplementary Note 3: Secure & Distributed Computation of a GWAS

Figure S2 depicts FAMHE secure and federated workflow for the computation of a Genome-Wide Association Study (see *Online Methods*). In Figure S2a, we describe FAMHE's protocol to compute an exact GWAS. First the covariance matrix ($X^T X$) is collectively computed (Collective Aggregation, CA) by the DPs before being inversed in the encrypted domain by one data provider ($DP_R$) in **Step 1**. In **Step 2**, the inversed matrix is augmented with the variant's contribution (vector $u$ that contains the variant's value for each patient) by following the Shermann-Morrison formula[48] and the method presented in the report on Cryptographic and Privacy-preserving primitives (page 52) of the WITDOM European project[49]. We remark that whenever possible, the DPs locally compute on their cleartext data before securely aggregating their partial results. The matrix $\boldsymbol{W}$ is made of 3 submatrices ($\boldsymbol{W}_{11}$, $\boldsymbol{W}_{12}$,$\boldsymbol{W}_{21}$) that are computed in Step 2. In **Step 3**, the model weights (or coefficients, $w$) are computed such that all elements required to obtain the p-value corresponding to the variant's coefficient can be computed in **Step 4**. In **Step 5**, the results are switched (Key Switching, KS) to the querier's public key such that she/he can compute the final result in **Step 6**. We remark that this process is performed simultaneously on multiple variants thanks to the SIMD property of the cryptoscheme, as described in Figure S3. In Protocol S2b, we show how the GWAS can be estimated and therefore computed with a lower complexity by avoiding the computation of the complete inverse matrix of the covariance matrix. The steps are similar as in Protocol S2a except that in **Step 2**, the model coefficients corresponding to the covariates are obtained through an efficient stochastic gradient descent with the label (phenotype) $y$. We rely on the protocols proposed by Froelicher et al.[25] to perform the SGD on an encrypted model. **Step 3** corresponds to a partial execution of steps 2 and 3 from Protocol S2a. In **Step 4**, we compute the coefficient corresponding to the variant and the other elements required to obtain the p-value.

**a** FAMHE-GWAS.

**Step 1: Inverse of covariance matrix of covariates**

Each $DP_i$: $X_i = [\mathbf{1}, X_i]$ with $X_i \in \mathbb{R}^{(p_i \times (f+1))}$

Each $DP_i$ computes $X_i^T X_i$ with $X_i^T X_i \in \mathbb{R}^{((f+1)\times(f+1))}$

CA $\rightarrow \boldsymbol{E(X^T X)}$

$DP_R$: $\boldsymbol{E((X^T X)^{(-1)}) = GJ(E((X^T X)))}$ and broadcasts

**Step 2: Variants contribution**

**For each** column $u$ in $V$ do:

Each $DP_i$ computes $u_i^T u_i$, $u_i^T X_i \times E((X^T X)^{(-1)})$

CA $\rightarrow \boldsymbol{E(u^T u), E(W_{21}) = E(u^T X (X^T X)^{(-1)})}$

$DP_R$ broadcasts $E(u^T X (X^T X)^{(-1)}))$

Each $DP_i$ computes $E(u^T X (X^T X)^{(-1)})) \times X_i^T u_i$

CA $\rightarrow \boldsymbol{E(u^T X (X^T X)^{(-1)} X^T u)}$

$DP_R$ computes $\boldsymbol{E(\frac{1}{c}) = E(u^T u - u^T X (X^T X)^{(-1)} X^T u)}$

Each $DP_i$ computes $E((X^T X)^{(-1)})) \times X_i^T u_i$

CA $\rightarrow \boldsymbol{E(W_{12}) = E((X^T X)^{(-1)} X^T u)}$

$DP_R$ computes $\boldsymbol{E(W_{11})}$=$\boldsymbol{E(\frac{1}{c}(X^T X)^{(-1)} + W_{12} W_{21})}$

$\boldsymbol{E(W) = E}\begin{pmatrix} W_{11} & W_{12} \\ W_{21} & 1 \end{pmatrix}$

**Step 3: All coefficients**

Each $DP_i$ computes $[X_i, u_i]^T y_i$

CA $\rightarrow \boldsymbol{E(w) = E(W \times [X, u]^T y)}$

**Step 4: P-value elements**

Each $DP_i$: $E(y_i') = X_i \times E(w)$ and $E(mse_i) = \frac{1}{p} \sum_{j=0}^{p_i} (E(y_i'[j]) - y_i[j])^2$

CA $\rightarrow E(mse)$

**Step 5: Key Switching**

$\mathrm{KS}_Q(E(mse), E(\frac{1}{c}, E(w[f+2]))$ and send to $Q$

**Step 6: Querier final result**

Querier decrypts and $p_{val} = 2 \cdot pnorm(-|\frac{w[f+2]}{\sqrt{(mse \cdot c)}}|)$

**b** FAMHE-FastGWAS.

**Step 1: Inverse of covariance matrix of covariates**

Each $DP_i$: $X_i = [\mathbf{1}, X_i]$ with $X_i \in \mathbb{R}^{(p_i \times (f+1))}$

Each $DP_i$ computes $X_i^T X_i$ with $X_i^T X_i \in \mathbb{R}^{((f+1)\times(f+1))}$

CA $\rightarrow \boldsymbol{E(X^T X)}$

$DP_R$: $\boldsymbol{E((X^T X)^{(-1)}) = GJ(E((X^T X)))}$ and broadcasts

**Step 2: Covariates coefficients & error**

$\boldsymbol{E(w[0:f+1])} = SGD(X, y)$ and broadcasts

Each $DP_i$: $E(y_i'') = y_i - E(X_i w[0:f+1])$

Each $DP_i$: $E(\bar{y_i''}) = \frac{1}{p} \sum_{j=0}^{p_i} E(y_i''[j])$

**Step 3: Variants contribution**

**For each:** column $u$ in $V$ do:

Each $DP_i$ computes $u_i^T u_i$, $u_i^T X_i \times E((X^T X)^{(-1)})$

CA $\rightarrow \boldsymbol{E(u^T u), E(W_{21}) = E(u^T X (X^T X)^{(-1)})}$

$DP_R$ broadcasts $E(u^T X (X^T X)^{(-1)}))$

Each $DP_i$ computes $E(u^T X (X^T X)^{(-1)})) \times X_i^T u_i$

CA $\rightarrow \boldsymbol{E(u^T X (X^T X)^{(-1)} X^T u)}$

**Step 4: P-value elements**

$DP_R$ compute $\boldsymbol{E(\frac{1}{c}) = E(u^T u - u^T X (X^T X)^{(-1)} X^T u)}$

Each $DP_i$: $E(\bar{u}_i) = \frac{1}{p} \sum_{j=0}^{p_i} E(u_i[j])$

CA $\rightarrow \boldsymbol{E(\bar{y''}), E(\bar{u})}$ and broadcasts

Each $DP_i$: $E(d_i) = \sum_{j=0}^{p_i} (u_i[j] - E(\bar{u}))^2$ and $E(t_i) = \sum_{j=0}^{p_i} (u_i[j] - E(\bar{u}))(y_i''[j] - E(\bar{y''}))$

CA $\rightarrow \boldsymbol{E(d), E(t)}$ and broadcasts

Each $DP_i$: $E(mse_i') = \sum_{j=0}^{p_i} (E(dy_i''[j]) - E(t) u[j])^2$

CA $\rightarrow \boldsymbol{E(mse')}$

**Step 5: Key Switching**

$\mathrm{KS}_Q(E(mse'), E(\frac{1}{c}), E(t), E(d)$ and send to $Q$

**Step 6: Querier final result**

Querier decrypts and $p_{val} = 2 \cdot pnorm(-|\frac{t/d}{\sqrt{\frac{1}{d^2}(mse' \cdot c)}}|)$

**Figure S2. Secure & Distributed Computation of a GWAS.** CA stands for collective aggregation, KS for collective key switching and bold indicates new values obtained from the DPs collaboration.

In Figure S3, we describe how the values are packed in order to perform the computations of the main variables used in the protocols presented in Figure S2. We note that 13 covariates are considered in the study that we reproduced[31]. We provide here the main intuition behind the packing of each element (in Figure S3) and its high level cost in terms of operations:

1. The inverse computation (line 4 in Protocol S2b & S2a) requires row operations on the input matrix. The matrix is therefore row-wise encrypted and its rows are duplicated to enable an efficient use of SIMD and to perform the subsequent operations for multiple variants simultaneously, thus minimizing rotations on encrypted data.

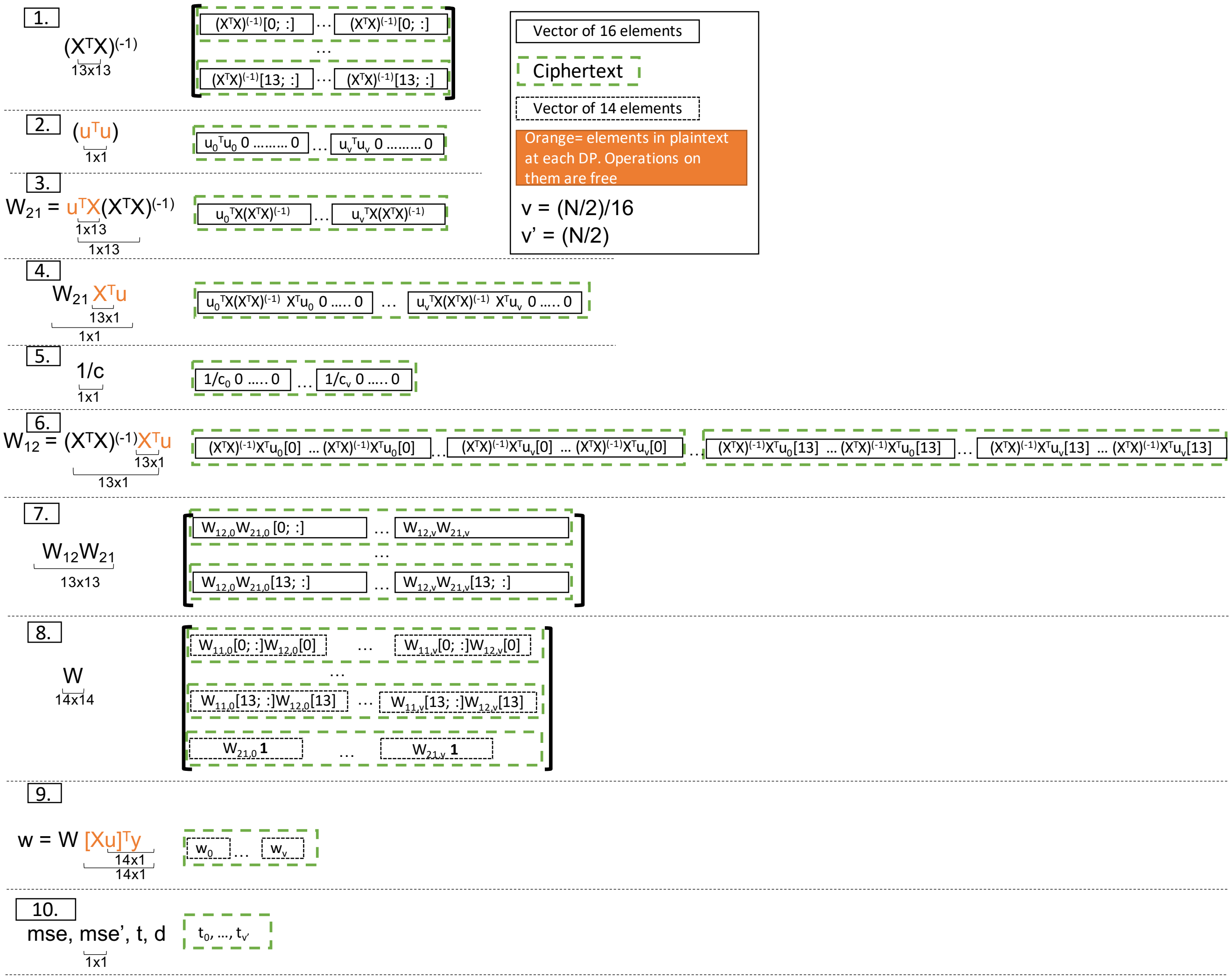


**Figure S3. Optimized Packing for a Simultaneous Computation on v Variants.**

2. The vectors multiplication results are packed such that they are in the right positions to perform the subtraction in line 11 for Protocol S2a and line 14 for Protocol S2b. This packing can be performed by the data providers on their cleartext data and does not require rotations of encrypted ciphertexts.

3. Each element of the plaintext $u^TX$ is replicated to be multiplied with a row of $(X^TX)^{(-1)}$ and the 13 multiplication results are aggregated. This operation does not require any rotation either.

4. The dot product between the plaintext vector $X^Tu$ and each encrypted row of $W_{21}$ is performed and the result is packed such that it can be used as such in the subsequent subtraction (line 11 for Protocol S2a and line 14 for Protocol S2b). The dot product requires $log_2(vectorsize)$ rotations. We note here that all vectors are padded with zeros such that their size is a power of 2. This is done to optimize the number of rotations required.

5. As before, the scalar values for $v$ variants are packed to simplify the subsequent subtraction.

6. As in 4, the dot product is performed between the plaintext vector $X^T u$ and each encrypted row of the matrix $(X^T X)^{(-1)}$. The results are then duplicated to prepare the multiplication of line 14 in Protocol S2a. This operation requires $2 \times log_2(vectorsize)$ rotations.

7. Each of the 13 ciphertexts of $W_{12}$ is multiplied with $W_{21}$. This operation does not require any rotation as the values have been prepared (packed) for this multiplication beforehand (in 6).

8. To construct the matrix $W$ for $v$ variants, we include $W_{12}$ directly in the padding of $W_{11}$. Due to $W_{12}$ packing, this is done in one mask (multiplication with binary vector) and one addition and does not require any rotation. Similarly, one addition is performed to include the 1 values in $W_{21}$.

9. The dot product between each encrypted row of $W$ and $[Xu]^T y$ is performed such that the result can be packed to prepare the multiplication of line 17 in Protocol S2a. This requires $log_2(vector/rowsize) + (vector/row - 1)$ rotations.

10. All elements are computed for $v$ variants simultaneously.

### Supplementary Note 4: Quantitative Comparison of FAMHE with existing approaches.

FAMHE is specifically designed to benefit from its multiparty construction and to optimize its use of MHE to efficiently execute secure FA workflows among a large number of data providers that keep their data locally. **A non-secure centralized solution** based on PLINK takes 14 seconds when performed by a single data provider on the pooled dataset. As shown in Figure 4a, FAMHE efficiently distributes its workload and achieves an execution time of 69 minutes when the data are split among 12 DPs for the same computation. **A non-secure federated solution** based on PLINK's meta-analysis method takes around 5 minutes when executed on 12 DPs but yields very imprecise results. We remark that in the meta-analysis the DPs exchange information only once at the end and a non-secure solution in which the DPs collaborate during the process would be slightly slower, e.g., between 5 and 10 minutes, depending on the communication settings. **Differential-privacy-based solutions** usually yield the same execution time as non-secure solutions. We estimate that a **centralized HE-based solution** would take at least 2228 (with the FastGWAS approach) and 30,781 minutes (with the GWAS approach), whereas FAMHE respectively takes 69 and 812 minutes when the same approaches are collaboratively executed by 12 data providers. The centralized approach, contrarily to FAMHE, cannot distribute the workload among multiple DPs and suffers from a high overhead brought by centralized cryptographic operations. For example, in a centralized setting, a ciphertext is refreshed or bootstrapped in 26 seconds[53] for a security level of 108 bits, whereas the corresponding interactive protocol in FAMHE takes 0.6 seconds with a better security level of 128 bits. **SMC approaches** usually target settings with 2 to 4 parties due to a communication cost that becomes prohibitive for higher numbers of parties. FAMHE also works with 2 to 4 parties but is not specifically optimized for this scenario and its execution time would be in the same order of magnitude as secret-sharing-based solutions. However, unlike secret-sharing-based solutions, FAMHE efficiently scales to federated learning settings where many DPs keep their data locally. Furthermore, by designing an alternative MHE-friendly algorithm for the same task (e.g. FAMHE-FastGWAS), FAMHE can achieve faster execution times than SMC even in the setting with a small number of parties.

We summarize this comparison in Table S2. The two *Decentralized* approaches refer to cleartext adaptations of FAMHE's approaches. The results for *Centralized, Non-Secure* and *Meta-analysis, Non-Secure* are obtained through experiments with the PLINK software. The execution times for the *Decentralized* approaches are inferred from *Meta-analysis, Non-Secure* and their communication overheads are deduced from FAMHE's communication costs from which we removed the encryption overhead. We also relied on this encryption space overhead to estimate *Centralized HE* communication cost, whereas its execution time is deducted from our previous observation on the centralized bootstrapping. We estimated the performance of SMC by extrapolating the published results of Cho et al.[19]. For a fair comparison, we considered only the Phase 3 results, i.e., the association tests, for which Cho et al. computed the $\chi^2$ statistics of the generalized Cochran-Armitage trend test corrected for covariates. We note that, while Cho et al. take a different approach to computing the association results than FAMHE, the overall workflow is similar and represents a suitable reference point for comparison. Since the prior work considered a three-party setting, we expect the real execution time of SMC to be higher in our setting with 12 DPs. For the communication cost of SMC, we combined the estimates of both initial data sharing and the main computation step (Phase 3) from the original publication and assumed linear scaling with the number of DPs. FAMHE measurements are obtained through our experiments.

| | Centralized, Non-Secure | Meta-analysis, Non-Secure | Decentralized Non-secure | Decentralized, DiffP. | Centralized HE | SMC | FAMHE |
|---|---|---|---|---|---|---|---|
| Execution Time (min) | 0.24 | 5 | 10 | 10 | 2228; 30781 | 320 | 69; 812 |
| Communication cost (GB) | 4 | 0.5 | 39; 171 | 39; 171 | 12 | 3144 | 122; 684 |

**Table S2. Quantitative comparison of existing approaches for biomedical FA.** In distributed cases, the data are split among 12 data providers. The communication cost corresponds to the amount of data that one DP has to send during the entire process. When two values are provided, the left one corresponds to an approach equivalent to FastGWAS and the right one to GWAS.

### Supplementary Note 5: Data Providers Independent Results

Figure S4 complements Figure 3e by showing the independent GWAS result obtained by each of the 12 data providers on its own subset of the data.

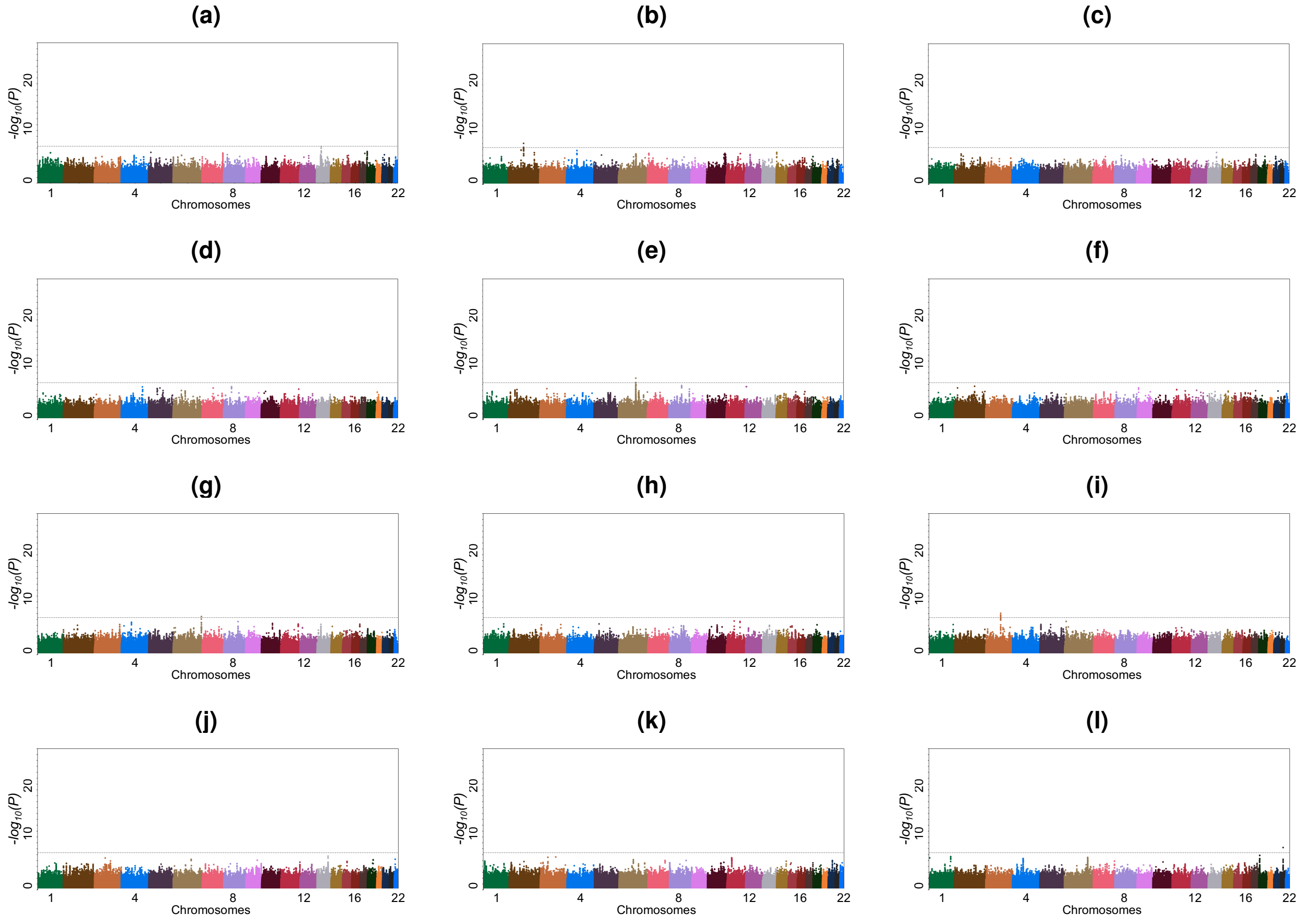


**Figure S4. Independent GWAS Results.** Results obtained by each data provider using only its own subset of the data.

### Supplementary Note 6: Table of Symbols

| Symbol | Description |
|---|---|
| FA | Federated Analytics |
| ML | Machine Learning |
| DP | Data Provider |
| DiffP | Differential Privacy |
| CA | Collective Aggregation |
| KS | Key Switching |
| MHE | Multiparty Homomorphic Encryption |
| SMC | Secure Multiparty Computation |
| $w$ | Model coefficients/weights |
| $pk$ | Collective public key |
| $N$ | Number of values encrypted in one ciphertext |
| SIMD | Single Instruction, Multiple Data |
| $S$ | Number of data providers |
| $t_j$ | Time when at least one event happened |
| $d_j$ | Number of events at time $t_j$ |
| $n_j$ | The number of individuals known to have survived (or at risk) |
| $\hat{S}(t)$ | Kaplan-Meier Estimator |
| $T$ | Number of data points |
| $p$ | Number of patients |
| $f$ | Number of features |
| $X \in \mathbb{R}^{p \times f}$ | Covariates matrix |
| $y \in \mathbb{R}^{p \times 1}$ | phenotype or label |
| $v$ | Number of variants |
| $u \in \mathbb{R}^{p \times 1}$ | Vector of 1 variant value for all patients |
| $V \in \mathbb{R}^{p \times v}$ | Variants matrix |
| $p_{val}$ | P-value |
| $pnorm$ | Cumulative distribution function (CDF) of the standard normal distribution |
| $w[f+2]$ | Weight corresponding to the variant |
| $y'$ | Prediction vector |
| $MSE$ | Mean Squared Error |
| $p_i$ | $DP_i$'s subset patients |
| $DP_R$ | Root of the tree |
| $E()$ | Encrypted value |
| $GJ$ | Gauss-Jordan method |
| $Q$ | Querier |

**Table S3. Frequently Used Symbols and Notations.**
